# Cyto-nuclear shuttling of afadin is required for rapid estradiol-mediated modifications of histone H3

**DOI:** 10.1101/284422

**Authors:** Katherine J. Sellers, Iain A. Watson, Deepak P. Srivastava

## Abstract

Estrogens have been shown to rapidly regulate local signalling events at both synapses and within the nucleus. The result of these signalling events is to rapidly modulate synapse structure and function, as well as epigenetic mechanisms including histone modifications. Ultimately these mechanisms are thought to contribute to long-lasting changes in neural circuitry, and thus influences cognitive functions such as learning and memory. However, the mechanisms by which estrogen-mediated local synaptic and nuclear signalling events are coordinated are not well understood. In this study we have found that the scaffold protein afadin, (also known as AF-6), undergoes a bi-directional trafficking to both synaptic and nuclear compartment in response to acute 17β-estradiol (estradiol) treatment. Interestingly, nuclear accumulation of afadin was coincidental with an increase in the phosphorylation of histone H3 at serine 10 (H3S10p). This histone modification is associated with the remodelling of chromatin into an open euchromatin state, allowing for transcriptional activation and related learning and memory processes. Critically, the cyto-nuclear trafficking of afadin was required for estradiol-dependent H3S10p. We further determined that nuclear accumulation of afadin is sufficient to induce phosphorylation of the mitogentic kinases ERK1/2 (pERK1/2) within the nucleus. Moreover, nuclear pERK1/2 was required for estradiol-dependent H3S10p. Taken together, we propose a model whereby estradiol incudes the bi-directional trafficking of afadin to synaptic and nuclear sub-compartments. Within the nucleus, afadin is required for increased pERK1/2 which in turn is required for H3S10p. Therefore this represents a mechanism through which estrogens may be able to coordinate both synaptic and nucleosomal events within the same neuronal population.

## Introduction

It is now well accepted that estrogens, and in particular 17β-estradiol (estradiol), can elicit rapid signaling actions in a range of cell types, including neurons (McEwen and Alves, 1999;Srivastava et al., 2013;Choleris et al., 2018). The rapid “non-conical” actions of estrogens are reliant on the activation of specific intracellular signaling pathways, and can influence a range of cellular events (Sellers et al., 2014;Choleris et al., 2018). This includes the remodeling of dendritic spines, the trafficking and post-translational modifications of proteins (Srivastava et al., 2008;Srivastava et al., 2010;Sellers et al., 2015), as well as transcriptional and epigenetic mechanisms (Boulware et al., 2005;Zhao et al., 2010;Zhao et al., 2012). Importantly, the rapid modulation of these non-conical mechanism have been shown to be crucial for the consolidation of new memories (Luine and Frankfurt, 2012;Srivastava et al., 2013;Choleris et al., 2018). Of these mechanisms, the regulation of epigenetic modifications is thought to be key in translating estradiol’s cellular effects into long-lasting influences on memory (Zhao et al., 2010;Zhao et al., 2012;Fortress and Frick, 2014).

There is accumulating evidence that post-translational modifications of histone proteins is a critical mechanisms for the remodeling of chromatin (Watson and Tsai, 2017). The phosphorylation or acetylation of histones is associated with the initiation of gene transcription (Berger, 2007), and are thought of as essential transcriptional regulatory mechanisms (Riccio, 2010;Maze et al., 2013;Watson and Tsai, 2017), which in turn have been shown to be required for long-lasting changes in behaviour. Interestingly, estradiol has been shown to cause acetylation of Histone H3 within 5 minutes in hippocampal neurons, which was necessary for estrogen-dependent consolidation of memory (Zhao et al., 2010;Zhao et al., 2012). Another modification important for memory and that can be induced in response to multiple extracellular stimuli is the phosphorylation of histone H3 protein at serine 10 on its N-terminal tail (Brami-Cherrier et al., 2007;Lubin and Sweatt, 2007;Wittmann et al., 2009;VanLeeuwen et al., 2014). Although such modifications are thought to be important for the encoding of long-lasting memories, the synaptic and cytoplasmic signalling cascades that connect local signalling at synapses with these nucleosomal events are not fully understood (Fainzilber et al., 2011). Furthermore, it is not fully understood whether estradiol regulates this epigenetic modification.

Recently it have become apparent that proteins residing at synaptic or cytoplasmic locations can translocate to the nucleus in response to specific stimuli (Jordan and Kreutz, 2009;Ch’ng and Martin, 2011;Fainzilber et al., 2011). It has been proposed that upon nuclear accumulation, that proteins that undergo cyto-nuclear translocation can participate in nuclear events that result in gene expression changes contributing to long-term alterations of synapses (Fainzilber et al., 2011;Ch’ng et al., 2012;Karpova et al., 2013). Indeed, growing evidence indicates that this is achieved through the modification of histone proteins. Recently, we have shown that the scaffold protein afadin, (also known as AF-6), bi-directionally traffics to nuclear and synaptic sub-compartments in response to activity-dependent stimulation (VanLeeuwen et al., 2014). Importantly, the accumulation of nuclear afadin was required for both long-lasting changes in synaptic morphology and time-dependent phosphorylation of histone H3 (VanLeeuwen et al., 2014). Interestingly, estradiol has been demonstrated to traffic afadin to synaptic locations (Srivastava et al., 2008). However, it is not known if estrogen also causes the trafficking of afadin to the nucleus, and additionally whether this action can influence the post-translational modification of histone H3. In the present study we have examined the nuclear content of afadin following acute exposure to estradiol. Furthermore, we have investigated whether estradiol can phosphorylate H3 at serine 10 (H3S10p), and whether this nucelosomal event requires afadin and the activation of the mitogenic kinase ERK1/2. Our data indicates that an afadin/ERK1/2 pathway is required for the rapid modulation of H3S10p by estradiol.

## Materials and Methods

### Reagents and plasmid constructs

The following antibodies were purchased: GFP mouse monoclonal (MAB3580: 1:1000), NeuN mouse monoclonal (clone A60; MAB377: 1:500), phospho-histone H3 serine 10 mouse (H3S10p) monoclonal (clone 3H10; 05-806: 1:200) and myc rabbit polyclonal (06-549: 1:500) were from Millipore; phospho-p44/42 MAPK (ERK1/2; Thr202/Tyr204) rabbit monoclonal (D13.14.4E; #4370: 1:1000/1:500), ERK1/2 mouse monoclonal (L34F12; #4696: 1:1000) were from Cell Signaling Technologies; GFP chicken polyclonal (ab13972: 1:10,000) and histone 3 (total) rabbit polyclonal (ab1791: 1:2000) were from Abcam; myc mouse monoclonal (9E10; Developmental Studies Hybridoma Bank, Iowa: 1:200); L/S-afadin rabbit polyclonal (AF-6; A0224: 1:1000/1:750), β-actin mouse monoclonal (clone AC-74; A5316: 1:1000) were from Sigma; DAPI was from Life Technologies. Estradiol (17β-estradiol) (E8875) was from Sigma; kinase inhibitor U0126 (MEK kinase inhibitor) (9903S) was from Cell Signaling Technologies. Plasmids used in this study were myc-l-afadin, and myc-afadin-NT which have been previously described (Xie et al., 2005).

### Neuronal culture and transfections

Mixed sex cortical neuronal cultures were prepared from Sprague-Dawley rat E18 embryos as described previously (Srivastava et al., 2011). Animals were habituated for 3 days before experimental procedures, which were carried out in accordance with the Home Office Animals (Scientific procedures) Act, United Kingdom, 1986. All animal experiments were given ethical approval by the ethics committee of King’s College London (United Kingdom). Cells were plated onto 18 mm glass coverslips (No 1.5; 0117580, Marienfeld-Superior GmbH & Co.), coated with poly-D-lysine (0.2mg/ml, Sigma), at a density of 3×10^5^/well equal to 857/mm^2^. Neurons were cultured in feeding media: neurobasal medium (21103049) supplemented with 2% B27 (17504044), 0.5 mM glutamine (25030024) and 1% penicillin:streptomycin (15070063) (all reagents from Life technologies). It should be noted that the neurobasal media contains phenol red, a compound with known estrogenic activity. Neuron cultures were maintained in presence of 200 µM D,L-amino-phosphonovalerate (D,L-APV, ab120004, Abcam) beginning on DIV (days *in vitro*) 4 in order to maintain neuronal health for long-term culturing and to reduce cell death due to excessive Ca^2+^ cytotoxicity via over-active NMDA receptors (Srivastava et al., 2011). We have previously shown that the presence or absence of APV in the culture media does not affect E2’s ability to increase spine linear density (Srivastava et al., 2008). Half media changes were performed twice weekly until desired age (DIV 23-25). The primary cortical neurons were transfected with the appropriate plasmid at DIV 23 for 2 days, using Lipofectamine 2000 (11668027, Life Technologies) (Srivastava et al., 2011). Briefly, 4-6 µg of plasmid DNA was mixed with Lipofectamine 2000 and incubated for 4-12 hours, before being replaced with fresh feeding media. Transfections were allowed to proceed for 2 days, after which cells were used for pharmacological treatment or ICC.

### Pharmacological treatments of neuron culture

All pharmacological treatments were performed in artificial cerebral spinal fluid (aCSF): (in mM) 125 NaCl, 2.5 KCL, 26.2 NaHCO3, 1 NaH2PO4, 11 glucose, 5 Hepes, 2.5 CaCl2, 1.25 MgCl2, and 0.2 APV). ACSF was used to ensured that there were no interactions between the pharmacological compounds being used and unknown elements within the culture media that may indirectly affect the experimental outcome. For kinase inhibitor experiments, neurons were pre-treated for 30 minutes before application of E2 directly over neurons. All compounds were dissolved in either 100 % ethanol or DMSO at a concentration of 10 or 1 mM, serially diluted to a 10X working concentration in aCSF, and applied directly to neuronal cultures. Final concentration of solvent was < 0.01%: vehicle control was made up of solvent lacking compound diluted as test compounds. Treatments were allowed to proceed for indicated time before being lysed for Western blotting or fixed for immunocytochemistry (ICC).

### Immunocytohistochemistry (ICC)

Neurons were fixed in either 4% formaldehyde/4% sucrose PBS for 10 minutes, or in 4% formaldehyde/4% sucrose PBS followed by 10 minutes fix with Methanol pre-chilled to - 20°C. Coverslips were then permeabilised and blocked simultaneously in PBS containing 2% normal goat serum and 0.2% Triton-X-100 for 1 hr at room temperature. Primary antibodies were added in PBS containing 2% normal goat serum for 2 hr at room temperature, or overnight at 4 °C, followed by 3 × 10 minutes washes in PBS. Secondary antibodies were incubated for 1 hr at room temp, also in 2% normal goat serum in PBS. Three further washes (15 minutes each) were performed before cells were incubated in DAPI if required. Finally, coverslips were mounted using ProLong antifade reagent (Life Technologies).

### Quantitative Analysis of Nuclear Immunofluorescence

Confocal images of double-stained neurons were acquired with a Leica SP-5 confocal microscope using a 63x oil-immersion objective (Leica, N.A. 1.4) as a z-series, or with a Zeiss Axio Imager Z1, equipped with an ApoTome using a 63x oil-immersion objective (Carl Zeiss, N.A. 1.4). Two-dimensional maximum projection reconstructions of images were generated and linear density calculated using ImageJ/Fiji (https://imagej.nih.gov/ij/) (Srivastava et al., 2011). For measurements of nuclear protein content, regions were drawn around nuclei, as delineated by NeuN or DAPI staining, and saved as regions of interest (ROI). These ROI were then applied to a corresponding image of afadin staining from which the mean average intensity was collected to determine nuclear immunoreactivity levels. Immunoreactivity levels of the nuclear localised proteins H3S10p or pERK1/2 were also measured within the nuclear ROI. Images were selected by examining a z stack series of images through the nucleus and choosing a central/representative plane.. Cultures directly compared were stained simultaneously and imaged with the same acquisition parameters. For each condition, 10-16 neurons from at least 3 separate experiments were used. Experiments were carried out blind to condition and on sister cultures.

### Quantitative Analysis of synaptic and dendritic Immunofluorescence

Synaptic and dendritic localization of afadin was quantified using MetaMorph (Srivastava et al., 2011). Images were acquired as described above. The background corresponding to areas without cells were subtracted to generate a “background-subtracted” image. Images were then thresholded equally to include clusters with intensity at least twofold above the adjacent dendrite. Regions along dendrites were outlined using the “Parameters” utility and the total grey value (immunofluorescence integrated intensity) of each cluster, or all clusters within a region were measured automatically. Quantification was performed as detailed above.

### Biochemistry cell fractionation

Subcellular fractions were prepared using the Proteo-Extract kit (EMD Biosciences) following the manufacturer’s recommendations. Lysates were subjected to Western blotting; membranes were probed with the appropriate antibodies.

### Statistical Analysis

All statistical analysis was performed in GraphPad. Differences in quantitative immunofluorescence, dendritic spine number were identified by Student’s unpaired t-tests. For comparisons between multiple conditions, the main effects and simple effects were probed by one-way-ANOVAs followed by Tukey Post Hoc analysis. Error bars represent standard errors of the mean. Between 3-5 independent cultures were used for all experiments.

## Results

### Acute Estradiol treatment causes bi-directional trafficking of Afadin

Previous studies have demonstrated that afadin is a dual residency protein present in both nuclear and cytosolic compartments, and specifically at synapses (Buchert et al., 2007;VanLeeuwen et al., 2014). Moreover, we have recently shown than activity-dependent signaling causes bi-directional trafficking of afadin to discrete nuclear and synaptic sites within the same cell (VanLeeuwen et al., 2014). As acute exposure to estradiol has been shown to also cause the enrichment of afadin at synapses (Srivastava et al., 2008), we reasoned that this may also cause afadin to traffic to the nucleus within the same cell. To test this, we treated primary cortical neurons with vehicle or 10 nM estradiol for 30 minutes and assessed afadin content within the nucleus and along dendrites in the same cell population. Following treatment with estradiol, afadin nuclear content was significantly higher than in vehicle treated cells (Figure 1 A and B). Afadin was congruently found to cluster along MAP2-positive dendrites (Figure 1 A and B). Interestingly, afadin was particularly enriched in puncta juxtaposed to MAP2-positive dendrites, indicative of an enrichment at synapses (Figure 1 A). We further validated these observations by performing Western blotting on membrane and nuclear fractions generated from treated neurons. Consistent with our immunocytochemical data, both long- and short-(l-, s-) afadin isoforms were increased in membrane and nuclear fractions following treatment with estradiol (Figure 1 C + D). Importantly, when we examined l/s-afadin levels in whole cell lysates, no difference in overall expression levels were found (Figure 1 E and F), suggesting that the clustering of l/s-afadin within membrane/synaptic and nuclear compartments following estradiol treatment was due to a redistribution of the protein as opposed to an increase in protein expression. Taken together, these data support a model where l/s-afadin is bi-directionally trafficked to nuclear and synaptic sites within the same cells following acute estradiol stimulation.

**Figure 1.**
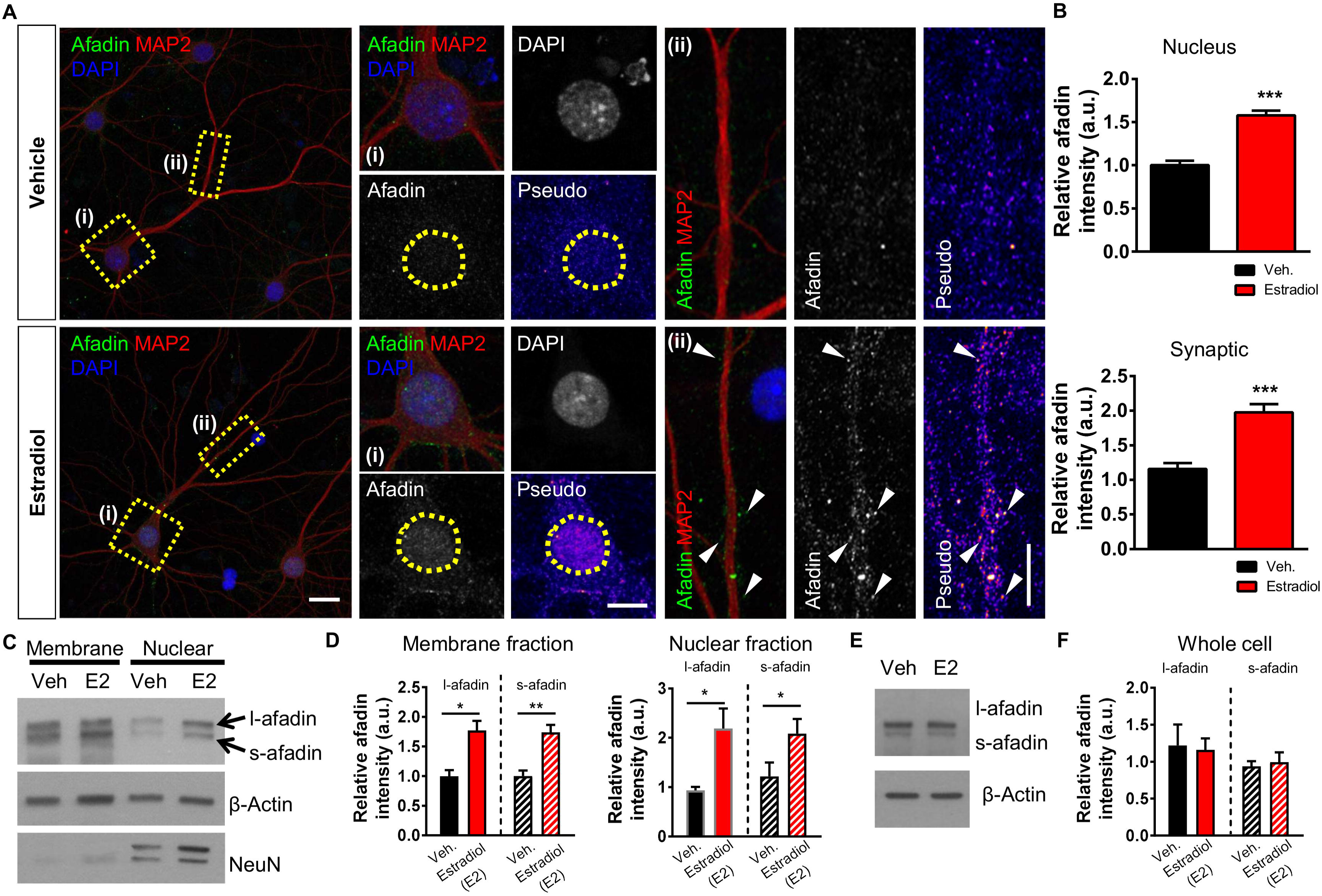
Estradiol induces the rapid bi-directional trafficking of afadin to synaptic and nuclear subcompartments. **(A)** Representative confocal images of DIV 23 cortical neurons treated with vehicle or 10 nM estradiol for 30 minutes. Insets show magnified regions of cell body including nucleus (‘**(i)**’, yellow box) (defined by DAPI staining) or dendrites (‘**(ii)**’, yellow rectangle) or within the same neuron. Pseudo images are of endogenous afadin content: yellow circles indicate the nucleus; white arrow heads indicate synaptic afadin puncta. **(B)** Quantification of afadin content at nuclear or synaptic subcompartments. Treatment with estradiol (30 minutes) causes a significant increase in afadin presence at both regions (***, p < 0.001, Student t-test; n = 80-81 cell per condition from 5 independent cultures). **(C)** Western blot analysis of membrane or nuclear cell fractions following treatment with vehicle or estradiol (also referred to as E2 in this figure). Blots were probed with an antibody that detects long-(l-) and short-(s-) afadin isoforms; β-actin was used a loading control and NeuN (Rbfox3) was used to confirm enrichment of nuclear compartment. **(D)** Quantification of l/s-afadin revealed an increase in both isoforms in membrane and nuclear fractions following treatment with estradiol (*, p < 0.05, **, p < 0.01, Student t-test; n = 3 independent cultures). **(E and F)** Assessment of afadin expression in whole cell following treatment with vehicle or estradiol reveals that there is no change in overall expression of the scaffold protein (Student t-test; n = 3 independent cultures). Scale bars = 20 µm, (i) 10 µm, (ii) 5 µm.

### Estradiol rapidly incudes phosphorylation of histone H3

The rapid post-translational modification of H3 has been observed in response to synaptic activity as well as neuromodulators (Brami-Cherrier et al., 2007;Stipanovich et al., 2008;Wittmann et al., 2009;VanLeeuwen et al., 2014). In particular, modulation of H3S10p by these stimuli, is thought to link extracellular signals with chromatin remodeling, and thus the regulation of genes associated with learning and memory (Riccio, 2010;Maze et al., 2013;Watson and Tsai, 2017). H3S10p is tightly regulated by mitogenic-kinases and is thought to be indicative of an open euchromatin state associated with transcriptional activation (Berger, 2007;Brami-Cherrier et al., 2009;Watson and Tsai, 2017). Interestingly, estradiol has been shown to modulate H3S10p via the Aurora kinase family to control cell proliferation in ovarian and tumor cells (Ruiz-Cortes et al., 2005;Mann et al., 2011). As estradiol has been shown to regulate a range of histone modification in neurons (Fortress and Frick, 2014), we were interested in understanding whether it also could rapidly regulate H3S10p levels. Remarkably, we observed a significant increase in H3S10p levels after just 30 minutes of treatment with estradiol (Figure 2 A and B). Next we investigated whether H3S10p was associated with the cytonuclear shuttling of afadin, as the presence of this scaffold protein in the nucleus is required for activity-dependent H3S10p (VanLeeuwen et al., 2014). Therefore we assessed whether estradiol caused a concomitant increase in both afadin and H3S10p levels within the same cells. When we limited our analysis only to neurons that demonstrated an increase in nuclear afadin compared to control, we still found a significant increase in H3S10p levels (Figure 2 C and D). Taken together, these data indicate that estradiol rapidly phosphorylates the N-terminal tail of H3 at serine 10, and that this occurs concurrently in neurons in which afadin traffics to the nucleus.

**Figure 2.**
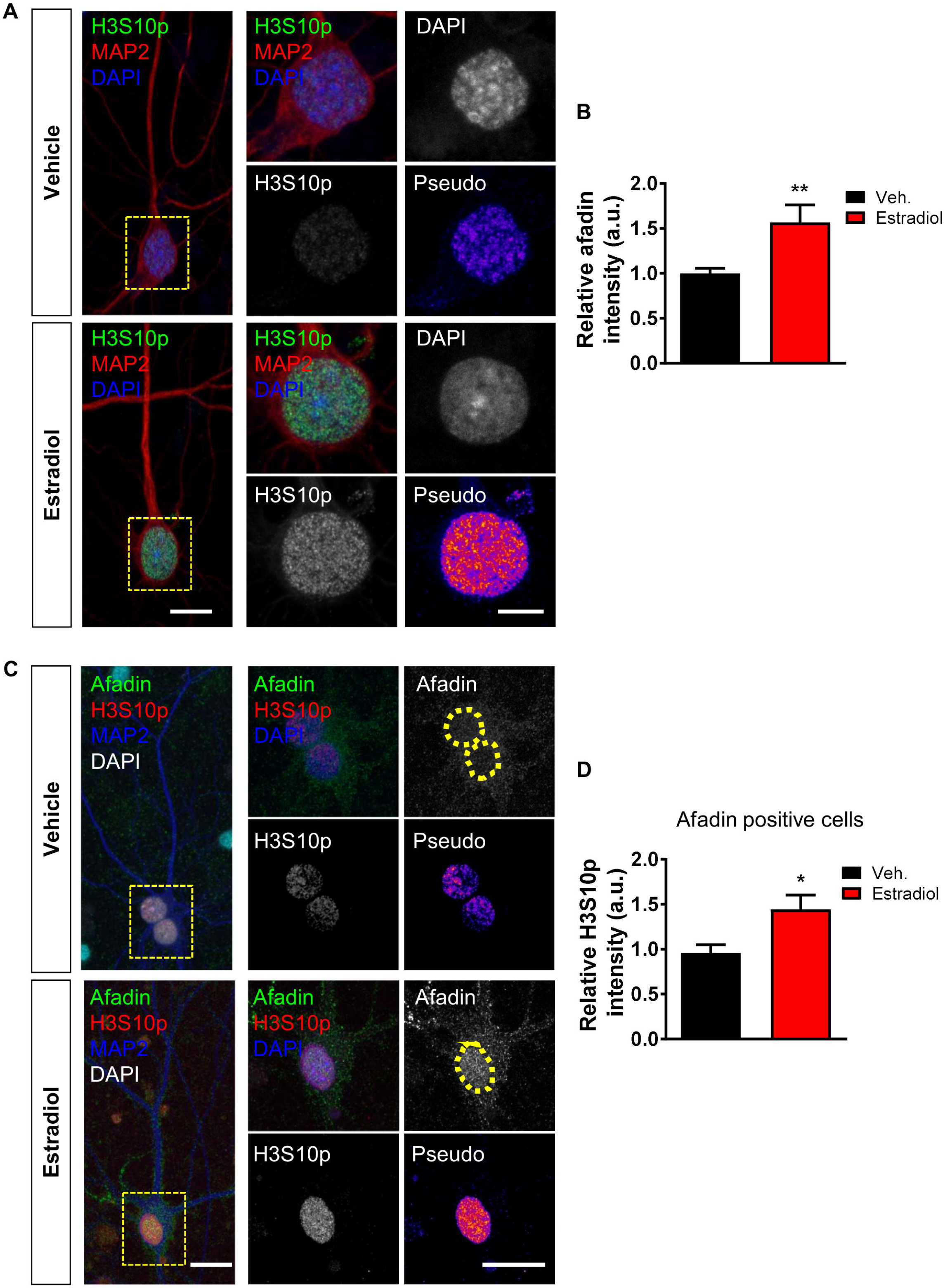
Estradiol-mediated histone H3 phosphorylation at serine 10 (H3S10p) and afadin nuclear accumulation occurs within the same population of neurons. **(A)** Representative confocal images of DIV 23 cortical neurons immunostained for MAP2 (morphological marker) and H3S10p; nucleus was identified by DAPI, after treatment with vehicle or estradiol (10 nM) for 30 minutes. Insets (yellow box) show magnified regions of cell body including nucleus defined by DAPI staining (yellow circle). Pseudo images are of endogenous H3S10p levels: ‘hot’ colours indicate increased expression. **(B)** Quantification of H3A10p levels reveals that estradiol treatment increases H3 phosphorylation (**, p <0.01, Student t-test; n = 47-48 cells per condition from 4 independent cultures). **(C)** Confocal images of DIV 24 cortical neurons treated with vehicle or estradiol and subsequently immunostained with afadin and H3S10p: DAPI was used to define nucleus. Insets (yellow box) show magnified regions of cell body including nucleus (yellow circle). Pseudo images are of endogenous H3S10p levels: ‘hot’ colours indicate increased expression. **(D)** Analysis of H3S10p levels in cells that exhibit increased afadin nuclear content. This analysis revealed that H3S10p is increased by estradiol in neurons that concurrently display an increase in afadin nuclear content (*, p <0.05, Student t-test; n = 25-28 cells per condition from 3 independent cultures). Scale bar = 10 µm; 5 µm.

### Estradiol-dependent phosphorylation of H3 is dependent on afadin

Our data above suggests a relationship between afadin trafficking to the nucleus and H3S10p. This is consistent with the requirement of increased nuclear afadin for activity-dependent H3S10p (VanLeeuwen et al., 2014). Therefore, to prevent the accumulation of afadin in the nucleus, we took advantage of the fact that the N-terminal region of afadin (afadin-NT) is required for the nuclear localization of the protein, and that exogenous expression of a myc-afadin-NT fragment blocks activity-dependent cytonuclear trafficking of endogenous afadin (VanLeeuwen et al., 2014). We first confirmed whether exogenous myc-afadin-NT could block estradiol-induced nuclear accumulation. Indeed we found that following treatment with estradiol, cells expressing myc-afadin-NT had an attenuated level of nuclear afadin compared to non-expressing cells (Figure 3 A and B). Interestingly, expression of the NT fragment did not alter levels of endogenous nuclear afadin under vehicle conditions (Figure 3 A and B), indicating that exogenous afadin-NT was only blocking the active transport of endogenous afadin into the nucleus. Next we treated neurons with estradiol or vehicle and examined whether exogenous myc-afadin-NT could block estradiol-induced H3S10p. Under basal conditions (Veh), exogenous afadin-NT had no alter H3S10p levels compared to non-expressing cells. However, following treatment with estradiol, cells expressing afadin-NT no longer demonstrated an increase in H3S10p levels (Figure 3 C and D). Taken together, these data suggest that the trafficking of afadin to the nucleus is required for estradiol-dependent phosphorylation of H3 at serine 10.

**Figure 3.**
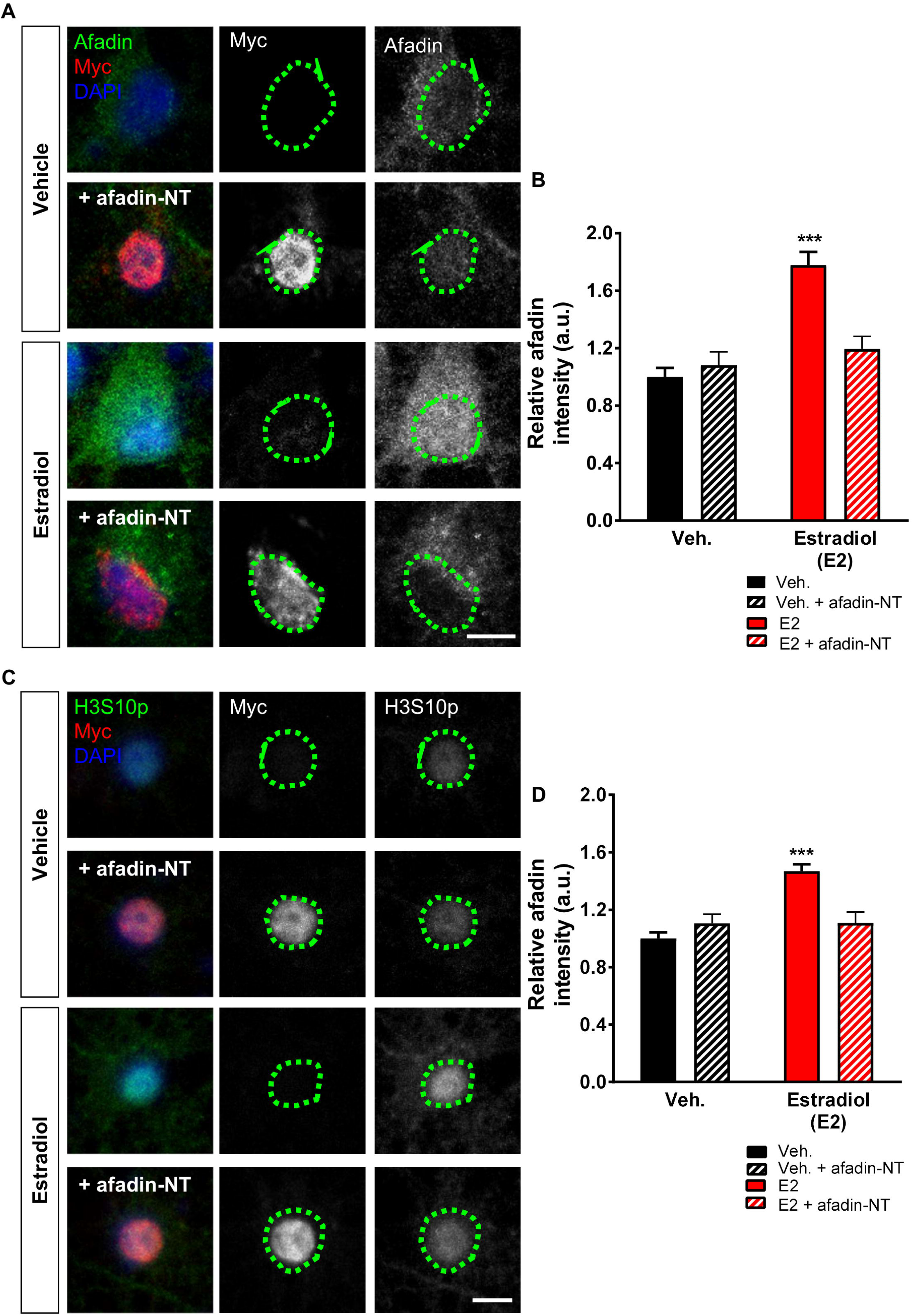
Cyto-nuclear trafficking of afadin is required for estradiol-induced increased H3S10p expression. **(A)** Representative confocal images of DIV 25 cortical neurons exogenously myc-afadin-NT, or not, treated with vehicle or estradiol (E2; 10 nM, 30 minutes), and subsequently fixed and immunostained for afadin and myc; nucleus was defined by DAPI (green circle). **(B)** Analysis of afadin nuclear content revealed that under basal conditions (vehicle treatment), afadin-NT did not alter afadin content in the nucleus. Conversely, afadin-NT attenuated estradiol-mediated afadin nuclear accumulation (F(3,41)=17.77, p < 0.001, Tukey Post Hoc, ***, p < 0.001, one-way ANOVA; n = 10-13 cells per condition from 3 independent cultures). **(C)** Confocal images of DIV 25 cortical neurons overexpressing myc-afadin-NT, or not, and treated with vehicle or estradiol (E2; 10 nM, 30 minutes). Neurons were fixed and subsequently immunostained for H3S10p and myc; nucleus was defined by DAPI (green circle). **(D)** Quantification of H3S10p levels following treatment revealed that exogenous afadin-NT reduced estradiol-mediated increased H3S10p expression (F(3,120)=19.11, p < 0.001, Tukey Post Hoc, ***, p < 0.001, one-way ANOVA; n = 20 – 50 cells per condition from 3 independent cultures). Scale bars = 5 µm.

### Acute estradiol treatment phosphorylates ERK1/2 in nucleus

One of the major kinases that mediates the rapid effects of estradiol is the mitogenic kinases, ERK1/2 (Sellers et al., 2014;Choleris et al., 2018). Interestingly, it has been reported that estradiol and specific estrogen receptor agonists can activate ERK1/2 via phosphorylation (pERK1/2) at multiple cellular compartments such as synapses and the nucleus (Mannella and Brinton, 2006;Srivastava et al., 2008;Srivastava et al., 2010). Consistent with this, we observed increased pERK1/2 levels along, and juxtaposed to dendrites, as well as within the nucleus after 30 minutes of estradiol treatment (Figure 4 A and B). To confirm these findings, we also probed nuclear fractions for pERK1/2 by Western blot. This further demonstrated that estradiol could rapidly increase pERK1/2 levels in the nucleus (Figure 4 C and D). Afadin is a component of the Rap/Ras signaling cascade, and therefore is thought to mediate Ras/Rap’s ability to signal to its downstream targets, which includes ERK1/2 (Ye and Carew, 2010;Woolfrey and Srivastava, 2016). We therefore reasoned that afadin may also be important for the activation of ERK1/2 in both dendritic and nuclear compartments. Indeed, overexpression of full length myc-l-afadin resulted in an increase in pERK1/2 levels (Figure E). Interestingly, pERK1/2 could be observed along dendrites, but also within the cell soma and nucleus (Figure 4 E). Taken together, these data support a model whereby estradiol increases ERK1/2 activation at multiple cellular compartments including the nucleus. Furthermore, afadin may play a role in mediating the subcompartment activation of ERK1/2.

**Figure 4.**
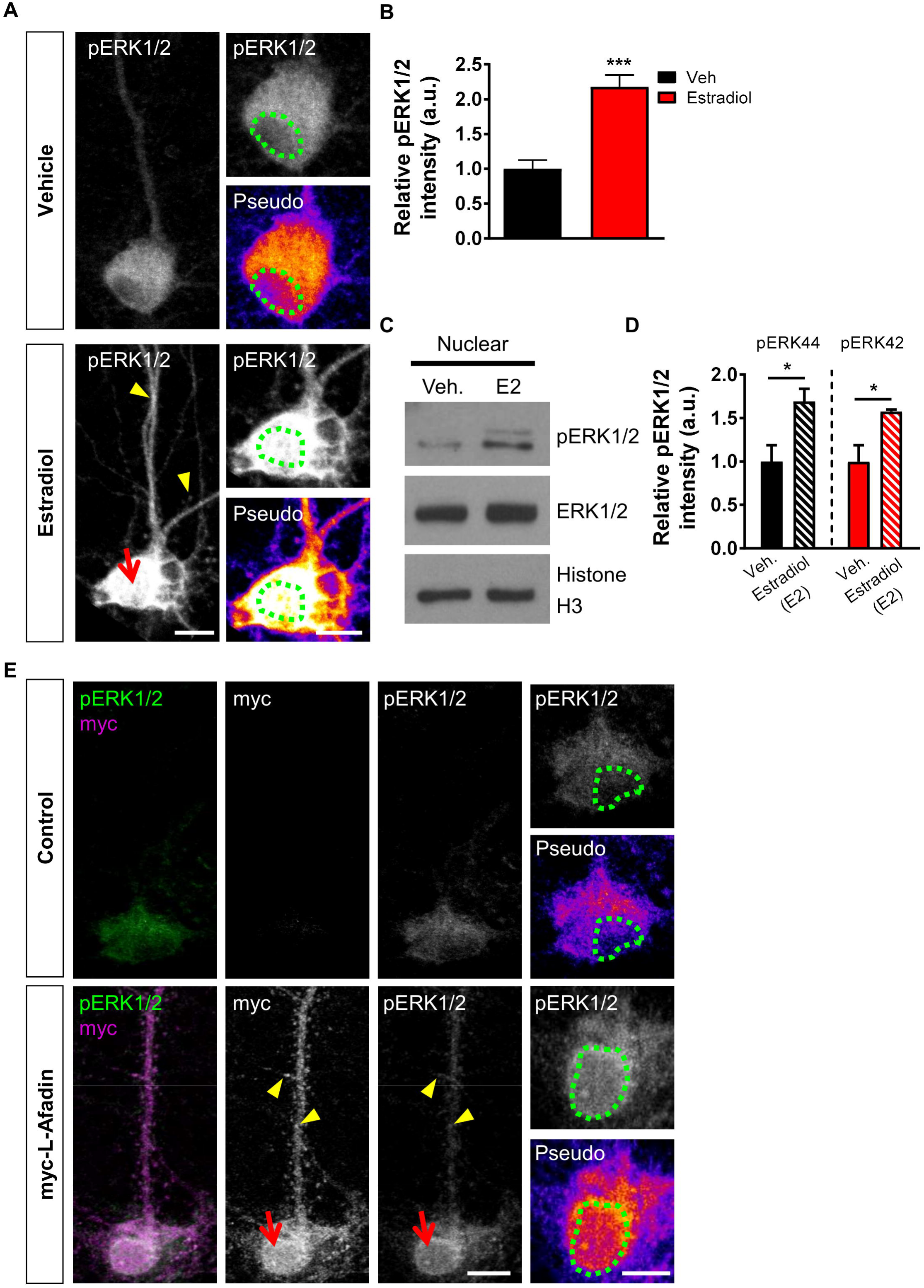
Estradiol rapidly phosphorylates nuclear ERK1/2. **(A)** Confocal images of DIV 25 cortical neurons treated with vehicle or estradiol (E2; 10 nM, 30 minutes), and subsequently fixed and immunostained for pERK1/2; nucleus was defined by DAPI (green circle). Pseudo images are of endogenous pERK1/2 levels: ‘hot’ colours indicate increased expression. **(B)** Quantification of pERK1/2 nuclear level revealed a significant increase in expression following estradiol treatment. (***, p < 0.001, Student t-test; n = 18-23 cell per condition from 3 independent cultures). **(C)** Western blotting of nuclear fraction of neurons treated with vehicle or estradiol (30 minutes): blots were probed for phospho and total ERK1/2; histone H3 was used to confirm enrichment of nuclear compartment. (D) Quantification revealed that estradiol caused a significant increase in pERK1/2 expression (*, p < 0.05,Student t-test; n = 3 independent cultures). (E) Representative confocal images of DIV 25 cortical neurons ectopically expressing myc-L-afadin or not and immunostained for myc or pERK1/2: nucleus was identified by DAPI. Pseudo images are of endogenous pERK1/2 levels: ‘hot’ colours indicate increased expression. Cells expressing myc-L-afadin demsontrated increased pERK1/2 levels along dendrites (yellow arrow heads) and within the nucleus. Scale bars = 10 µm or 5 µm.

### Estradiol-induced H3S10p requires ERK1/2 phosphorylation

As estradiol rapidly increased pERK1/2 levels in the nucleus, we reasoned that this kinase may be part of the pathway resulting in increased H3S10p levels. Consistent with this, the mitogenic kinases MSK1 and RSK1, which directly phosphorylate H3 at serine 10, are direct targets of ERK1/2 (Chwang et al., 2006;Brami-Cherrier et al., 2009;Ciccarelli and Giustetto, 2014;Bluthgen et al., 2017). Therefore, we tested whether estradiol-mediated H3S10p could be blocked by inhibiting pERK1/2 using the MEK kinase inhibitor U0126. Pretreatment with U0126 alone had no effect on basal levels of H3S10p levels (Figure 5A and B). However, pretreatment with U0126 significantly attenuated the ability of estradiol to increase H3S10p levels (Figure 5 A and B). Collectivity these data indicate that estradiol can rapidly increase H3S10p via an ERK1/2-dependent pathway.

**Figure 5.**
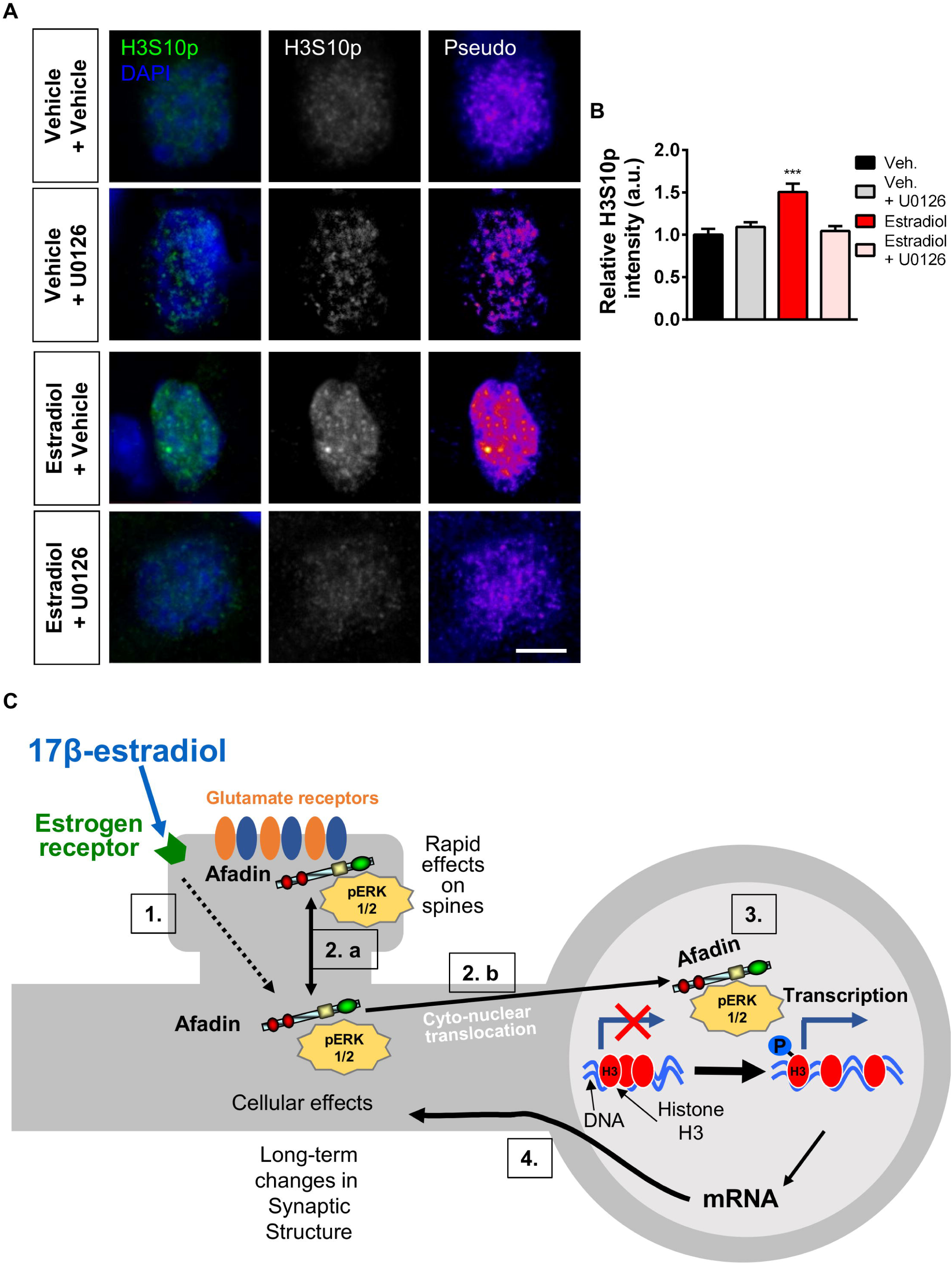
ERK1/2 activation is required for estradiol-dependent increased H3S10p expression. **(A)** Confocal images of DIV 23 cortical neurons pre-treated with MEK inhibitor, U0126 (10 µM for 20 minutes) or control, before being treated with vehicle or estradiol (E2; 10 nM, 30 minutes), and subsequently fixed and immunostained for H3S10p; nucleus was defined by DAPI (green circle). Pseudo images are of endogenous H3S10p levels: ‘hot’ colours indicate increased expression. **(B)** Quantification revealed that pre-incubation with U0126 blocked estradiol-mediated increased H3S10p expression (F(3,104)=10.01, p < 0.001, Tukey Post Hoc, ***, p < 0.001, one-way ANOVA, n = 24-29 cell per condition from 3 independent cultures). **(C)** Model of estradiol’s ability to coordinate synaptic and nuclear signalling. Scale bar = 5 µm.

## Discussion

In this study we have demonstrated that estradiol causes the bi-directional trafficking of afadin to synaptic and nuclear compartments within the same cell. Furthermore, we present evidence that estradiol phosphorylates histone H3 at serine 10, which is dependent on the nuclear accumulation of afadin. Interestingly, we also found that estradiol caused an increase in pERK1/2 levels within the nucleus, which could also be mimicked by the overexpression of l-afadin, and moreover, that the MEK/ERK1/2 signaling cascade was required for estradiol-dependent increases in H3S10p levels. Taken together, we propose a model where afadin is bi-directional trafficked to synaptic/membrane regions and the nucleus, where it is required for nuclear pERK1/2 and H3S10p increases in response to estradiol (Figure 5 C). This pathway may therefore represent a novel mechanism by which estrogens can rapidly remodel chromatin structure, and potentially regulate gene expression in a non-canonical manner.

The molecular mechanisms that enable neurons to translate signals generated at dendrites and synapses to the nucleus in order to regulate nucleosomal events are not well understood (Cohen and Greenberg, 2008;Jordan and Kreutz, 2009;Ch’ng and Martin, 2011). One mechanism by which signals initiated at the synapse can be propagated to the nucleus is through the propagation of by calcium signalling (Cohen and Greenberg, 2008). However, there is now growing evidence that the cyto/synto-nuclear shuttling of proteins provides another mechanism of signalling to the nucleus. (Jordan and Kreutz, 2009;Ch’ng and Martin, 2011;Fainzilber et al., 2011). Although this signalling modality is slower than the propagation of calcium signals to the nucleus, it potentially offers greater specificity and temporal regulation. Indeed, several proteins that display dual localization in the cytosol and nucleus of pyramidal neurons translocate to the nucleus following activation of NMDARs (Abe and Takeichi, 2007;Jordan et al., 2007;Proepper et al., 2007;Dieterich et al., 2008;Ch’ng et al., 2012;Karpova et al., 2013;VanLeeuwen et al., 2014). Once within the nucleus, these proteins may mediate gene transcription through the recruitment of different transcription factors, or indeed regulate post-translational modifications of histone proteins. Interestingly, this modality may also offer a level of temporal regulation that may not be possible by relying solely on calcium signalling. For example, we have previously shown that afadin shuttles to the nucleus and regulates H3S10p levels via a p90RSK-dependent mechanism in response to activity-dependent stimulation. However, this pathway was only required for the phosphorylation of H3 after 120 minutes of stimulation, and not at earlier time points (VanLeeuwen et al., 2014).

The data we presented in this study above is consistent with our previous work whereby following stimulation, afadin is trafficked to synaptic/membrane subcompartments, but also shuttles to the nucleus within the same cell (Srivastava et al., 2008;VanLeeuwen et al., 2014). We have proposed that this bi-directional trafficking of afadin potentially arises from two separate pools of protein. These distinct populations allow this protein to engage with local signalling events at synapses in addition to regulation of nucleosomal events (Figure 5 C). Consistent with this idea, we have previously shown that afadin is required for the maintenance of synaptic structure as well as excitatory tone (Srivastava et al., 2012). Critically, afadin is also necessary for the rapid remodelling, within 30 minutes, of dendritic spines in response to estradiol (Srivastava et al., 2008). Interestingly, within this time frame the remodelling of spines by estrogens occur independently of protein synthesis, and thus are not reliant on gene transcription (Srivastava et al., 2008). However, the cytonuclear shuttling of afadin may influence gene transcription, although how and whether this results in long-lasting changes in synaptic structure or function is currently unclear. Therefore, the increased presence of afadin at synaptic/membrane and nuclear compartment are likely to play different roles.

Multiple studies have shown the that histone H3 is phosphorylated in response to a variety of stimuli. Importantly, this modification has been established to be sufficient to confer a change in chromatin from a condensed heterochromatin state to a euchromatin state; more amenable to gene transcription and associated transcriptional activation (Berger, 2007;Maze et al., 2013;Watson and Tsai, 2017). H3S10p can be induced by activity-dependent stimuli, multiple neurotransmitters, learning and in response to drugs of abuse (Kumar et al., 2005;Chwang et al., 2006;Brami-Cherrier et al., 2007;Stipanovich et al., 2008;Brami-Cherrier et al., 2009;Wittmann et al., 2009;VanLeeuwen et al., 2014). Regulation of such nucleosomal events likely controls global and long-lasting changes in gene transcription, which may ultimately result in established changes to neuronal structure, and by extension synaptic function.

Previous studies have demonstrated that estrogens can regulate H3S10p to control cell proliferation in tumour and ovarian cells (Ruiz-Cortes et al., 2005;Mann et al., 2011). However, in the CNS estrogens have only been shown to regulate the acetylation of H3 in the CA1 region of the hippocampus; a requirement for estrogen-dependent memory consolidation (Zhao et al., 2010;Zhao et al., 2012). Interestingly, H3S10p is thought to be able to regulate gene transcription directly, but also to facilitate the acetylation of H3 (Lo et al., 2000;Riccio, 2010;Maze et al., 2013;Watson and Tsai, 2017). Future studies will be aimed at disentangling the potential relationship between these 2 different histone modifications following treatment with estradiol. Indeed, a critical question is whether H3S10p is required for estradiol-dependent memory consolidation. Parenthetically, previous studies have shown that estradiol also increases the phosphorylation of the transcription factor ‘cyclic AMP response element-binding protein (CREB) (Boulware et al., 2005). It would therefore be interesting to determine whether H3S10p and pCREB induced by acute estradiol exposure, act in concert or independently.

A surprising finding in this study, is the potential link between afadin, ERK/2 and H3S10p. Previous studies have shown that estradiol rapidly increases pERK1/2 levels in the nucleus (Mannella and Brinton, 2006), and moreover the pERK1/2 is required for estradiol-dependent increases in pCREB (Boulware et al., 2005). We find that in addition to estradiol treatment, overexpression of afadin also increased pERK1/2 levels within the nucleus. Critically, increased pERK1/2 levels were also seen along dendrites, consistent with the spatial distribution of pERK1/2 seen following activation of specific estrogen receptors (ERs) (Srivastava et al., 2010). At synaptic sites, afadin has been shown to act as a scaffold protein, allowing components of signalling pathways to interact (Xie et al., 2005;Srivastava et al., 2008;Xie et al., 2008;Srivastava et al., 2012). Building on this, we have previously suggested that one additional function that afadin may have is to mediate the transport of signalling proteins to the nucleus, and/or to act as a scaffold to assemble transcription factors or histone modifying proteins at discrete nuclear subcompartments (VanLeeuwen et al., 2014). Of note, uncaging of glutamate at a subset of spines is sufficient to cause the translocation of pERK1/2 to the nucleus (Zhai et al., 2013). In addition to translocating to the nucleus, pERK1/2 is required for H3S10p in response to multiple stimuli (Chwang et al., 2006;Brami-Cherrier et al., 2009;Ciccarelli and Giustetto, 2014), and also required for activity-dependent transcriptional regulation (Bluthgen et al., 2017). Therefore, one may propose a model whereby following acute estradiol treatment, pERK1/2 and afadin traffic to the nucleus as a complex, whereupon they increase H3S10p levels (Figure 5 C). Future studies will be aimed at determining the nature of the relation between these two proteins.

Taken together, this study provides the first evidence that estradiol can phosphorylate histone h3 in neurons. More specifically estradiol induces the cytonuclear trafficking of afadin and subsequent nuclear ERK1/2 activation of H3S10p under stimulated conditions. These data are consistent with growing studies that demonstrate that estrogens are capable of modulating gene transcription and protein synthesis in a non-conical manner (Srivastava et al., 2013;Sellers et al., 2014;Choleris et al., 2018). Moreover, we further provide evidence that estradiol can promote the bi-directional trafficking of afadin to both synaptic and nuclear compartments; this may underlie how estradiol exerts rapid, as well as long-lasting effects on synaptic structure and function. Significantly, understanding the downstream targets or gene modulated by this pathway, may reveal how estrogenic-modulation of this cytonuclear pathway may result in long-lasting changes to synaptic function.

## Acknowledgments

This work was supported by grants from Medical Research Council, MR/L021064/1, Royal Society UK (Grant RG130856), the Brain and Behavior Foundation (formally National Alliance for Research on Schizophrenia and Depression (NARSAD); Grant No. 25957), awarded to D.P.S.; K.J.S. was supported by a McGergor Fellowship from the Psychiatric Research Trust (Grant McGregor 97) awarded to D.P.S.; I.A.W. is funded by an ARUK studentship grant no. PhD2016-4, awarded to D.P.S.

## Author Contributions Statement

K.J.S, I.A.W. and D.P.S. conducted and analysed experiments; D.P.S. designed and conceptualized the study; K.J.S, I.A.W. and D.P.S. contributed to the writing; D.P.S. oversaw the project.

